# Human cytomegalovirus blocks canonical TGFβ signaling during lytic infection to limit induction of type I interferons

**DOI:** 10.1101/2021.02.16.431411

**Authors:** Andrew H. Pham, Jennifer Mitchell, Sara Botto, Kara M. Pryke, Victor R. Defilippis, Meaghan H. Hancock

## Abstract

Human cytomegalovirus (HCMV) microRNAs (miRNAs) significantly rewire host signaling pathways to support the viral lifecycle and regulate host cell responses. Here we show that SMAD3 expression is regulated by HCMV miR-UL22A and contributes to the IRF7-mediated induction of type I IFNs and IFN-stimulated genes (ISGs) in human fibroblasts. Addition of exogenous TGFβ interferes with the replication of a miR-UL22A mutant virus in a SMAD3-dependent manner in wild type fibroblasts, but not in cells lacking IRF7, indicating that downregulation of SMAD3 expression to limit IFN induction is important for efficient lytic replication. These findings uncover a novel interplay between SMAD3 and innate immunity during HCMV infection and highlight the role of viral miRNAs in modulating these responses.

**Author Summary:** Cells trigger the interferon (IFN) response to induce the expression of cellular genes that limit virus replication. In turn, viruses have evolved numerous countermeasures to avoid the effects of IFN signaling. Using a microRNA (miRNA) mutant virus we have uncovered a novel means of regulating the IFN response during human cytomegalovirus (HCMV) infection. Lytic HCMV infection induces the production of TGFβ, which binds to the TGFβ receptor and activates the receptor-associated SMAD SMAD3. SMAD3, together with IRF7, induces the expression of IFNβ and downstream IFN-stimulated genes in human fibroblasts. To counteract this, HCMV miR-UL22A, along with other HCMV gene products, directly targets SMAD3 for downregulation. Infection of fibroblasts with a miR-UL22A mutant virus results in enhanced type I IFN production in a SMAD3-dependent manner and the virus is impaired for growth in the presence of TGFβ, but only when both SMAD3 and IRF7 are present, highlighting the unique interaction between TGFβ and innate immune signaling.

## Introduction

Human Cytomegalovirus (HCMV) has co-evolved with its host over millions of years, resulting in exquisite control of both the cellular environment and the viral lifecycle that is highly cell type-dependent. Successful viral gene expression depends on blocking the powerful innate antiviral responses induced by viral binding and entry to limit interferon (IFN) production and signaling that act to render cells less permissive to viral infection (1). To evade the innate immune response, HCMV encodes numerous gene products that block the induction of intrinsic antiviral responses and the production of IFN and IFN stimulated genes (ISGs) (reviewed in (2, 3)). Along with IFNs, viral infection results in the secretion of additional cellular and viral cytokines and chemokines, which act in an autocrine and paracrine fashion to alter the intracellular and extracellular environment (4). For example, lytically infected cells (5-7) as well as latently-infected CD34^+^ hematopoietic progenitor cells (HPCs) (8-10), produce and secrete TGFβ which causes myelosuppression and has significant implications for hematopoietic stem cell transplantation (9).

TGFβ is a powerful regulator of numerous cellular pathways and has important roles in inflammation, immune modulation and cellular differentiation. The specific transcriptional outcomes of TGFβ signaling depends on cell type, TGFβ concentration and presence of additional signaling regulators resulting in either transcriptional activation or repression of different subsets of cellular genes (reviewed in (11, 12)). The complex and context-dependent outcome of canonical TGFβ signaling initiates with a relatively simple signal transduction pathway. TGFβ binding results in assembly of the receptor complex consisting of TGFβ receptors I and II. Receptor-associated SMADs (R-SMADs) SMAD2 and/or SMAD3 are then recruited and phosphorylated. Phosphorylation of R-SMADs enhances their interaction with the co-SMAD SMAD4 which results in shuttling of the R-SMAD/SMAD4 complex to the nucleus. SMADs regulate transcription by altering chromatin structure and generally have only weak affinity for the SMAD binding element (CAGAC) (13-15), therefore other DNA binding proteins are required for selective binding of the SMAD complexes to target elements. In addition to the choice of DNA binding partners, the recruitment of transcriptional coactivators, such as CBP/p300 (16) or corepressors, such as TGIF (17), SKI or SnoN (18) is also critical for determining the outcome of TGFβ signaling. Thus, the transcriptional outcome of canonical TGFβ signaling critically depends on the presence of a SMAD DNA binding cofactor as well as coactivators or corepressors, whose expression and localization are regulated by additional cellular signaling pathways in a context-dependent and cell type-specific manner (12).

Given the critical role that TGFβ plays in many cellular functions, activation and signaling by TGFβ is carefully regulated by the cell. One of the more recently studied means of regulation of the TGFβ signaling pathway is through the expression of cellular miRNAs (19). miRNAs are small, ∼22 nucleotide regulatory RNAs that post-transcriptionally regulate expression of genes through binding regions of complementarity in the targeted transcript. Sequence recognition occurs through the ‘seed region’ of miRNAs, nucleotides 2-8, and most commonly the 3’ UTR of the targeted gene, although binding to other regions of the transcript can also mediate regulation (20). miRNAs are the mRNA recognition component of a larger multi-protein RNA-induced silencing complex (RISC) that recruits proteins that mediate translational repression and/or mRNA degradation to the targeted transcript (21). Cellular miRNAs regulate the TGFβ signaling pathway at every level. Ligands, receptors, R-, inhibitory (I)-and co-SMADs and non-SMAD pathway components are all targets of cellular miRNAs (19). By targeting components of the signaling pathway as well as downstream transcriptional targets, miRNAs regulate all aspects of the intricate negative and positive feedback networks of the TGFβ signaling pathway.

HCMV also encodes its own miRNAs (22), and we have previously demonstrated that HCMV miR-UL22A-5p and −3p block canonical TGFβ signaling in latently-infected CD34^+^ HPCs by decreasing expression of the critical R-SMAD SMAD3 (9). Importantly, infection with a ∆miR-UL22A mutant virus restored TGFβ signaling to levels observed in mock infected HPCs and the mutant virus was impaired for reactivation from latency due to a loss of viral genomes or viral genome-containing cells. The attenuation of canonical TGFβ signaling in CD34^+^ HPCs as well as maintenance of the viral genomes during latency was restored by expression of a SMAD3 shRNA from the ∆miR-UL22A genome, indicating that targeting SMAD3 and the canonical TGFβ signaling pathway is essential to maintain the viral genome in latently infected cells (9). The mechanism of TGFβ-mediated viral genome loss remains to be determined and the effect of TGFβ signaling on other stages of the HCMV lifecycle is also still unknown.

Here we show that canonical TGFβ signaling negatively affects lytic replication of HCMV and is counteracted by reduced SMAD3 expression mediated, in part, by miR-UL22A. We show that miR-UL22A targeting of SMAD3 is critical for viral replication in the presence of TGFβ through attenuating IRF7-mediated activation of type I IFNs and ISGs. This study uncovers a novel link between SMAD3 and IRF7 in the induction of type I IFNs, highlighting the crosstalk between TGFβ and innate immune signaling during HCMV infection.

## Results

### Canonical TGFβ signaling is impaired during lytic HCMV infection

Our work (9) and that of others (6-8, 23, 24) has shown that CMV-infected cells produce and secrete TGFβ. We hypothesized that HCMV protects the lytically infected cell from the effects of TGFβ signaling by manipulating components of TGFβ signaling pathway, as we have observed during latent infection in CD34^+^ HPCs (9). To test this hypothesis, we infected normal human dermal fibroblasts (NHDF) (Fig 1A, C) and primary human aortic endothelial cells (hAEC) (Fig 1B, D) with the wild type TB40/E strain of HCMV for 48 hours followed by overnight serum starvation and 4 hour stimulation with TGFβ. In contrast to mock-infected cells, which respond to TGFβ treatment by upregulating the TGFβ-responsive transcripts JunB and SERPINE, HCMV-infected fibroblasts and endothelial cells do not show an enhancement in transcript expression following TGFβ treatment (Fig 1A, B). We next harvested protein lysates from mock-and HCMV-infected cells treated with TGFβ and analyzed expression of total and phosphorylated levels of SMAD3. Levels of both total and phosphorylated SMAD3 are reduced in HCMV-infected fibroblasts (Fig 1C) and endothelial cells (Fig 1D) compared to mock-infected cells. These data indicate that lytic HCMV infection reduces the levels of SMAD3 protein which contributes to a block in canonical TGFβ signaling.

**Fig 1.**
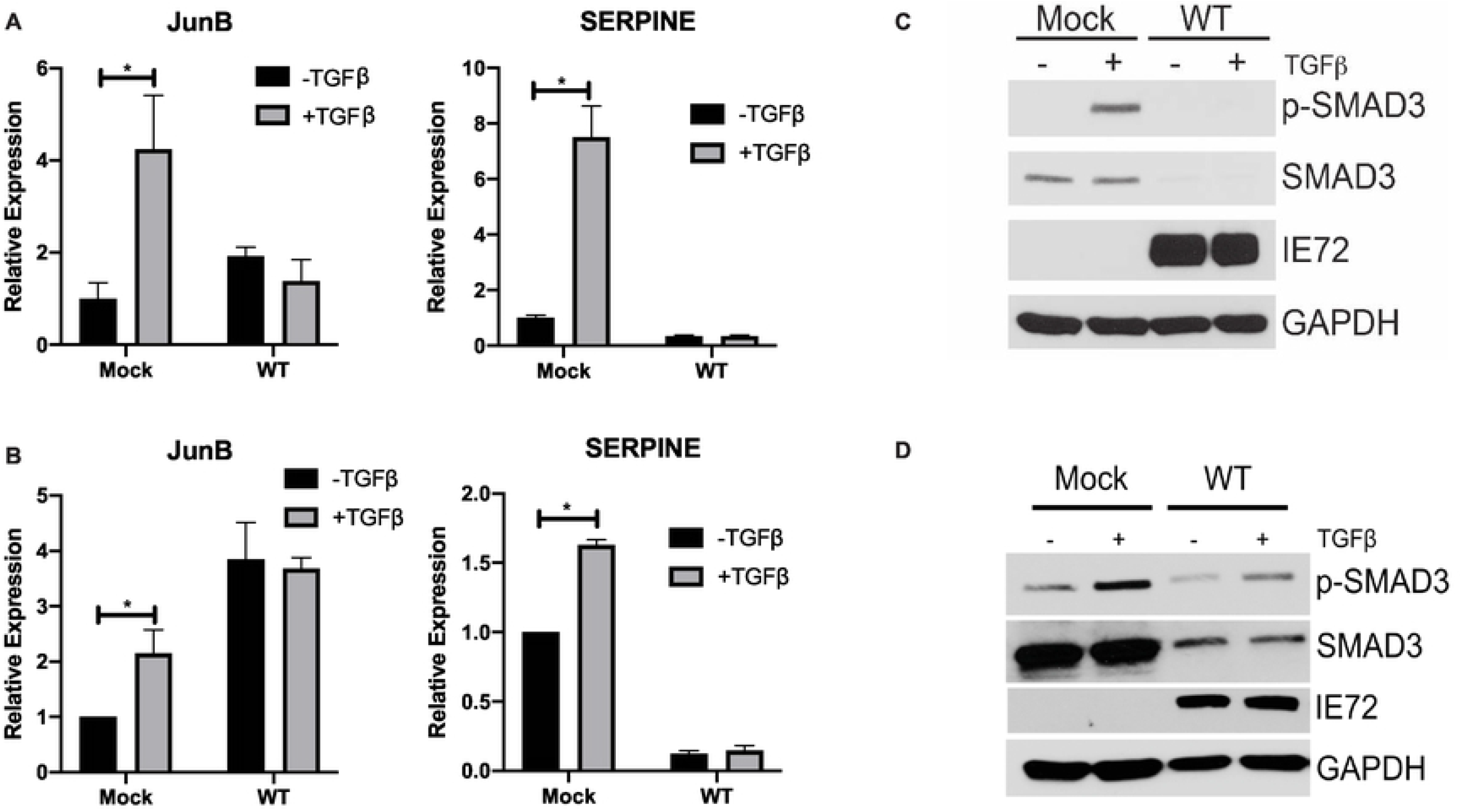
HCMV infection blocks canonical TGFβ signaling. NHDF (A) of hAEC (B) were infected with WT TB40E for 48 hours, followed by overnight serum starvation and stimulation with recombinant TGFβ (100pg/mL) for 4 hours. RNA was isolated and subjected to qRT-PCR for JunB and SERPINE. Experiments were performed in triplicate. * p<0.05 by two tailed Student’s t test. NHDF (C) or hAEC (D) were infected as above and protein lysates subjected to immunoblotting using the indicated antibodies.

### Mutation of miR-UL22A partially relieves the block to canonical TGFβ signaling observed during HCMV infection

We have previously identified SMAD3 as a target of the HCMV miRNAs miR-UL22A-5p and −3p, and have shown that miR-UL22A-mediated reduction of SMAD3 is required for genome maintenance during latent infection (9). To investigate the contribution of miR-UL22A targeting of SMAD3 to the block in canonical TGFβ signaling observed during HCMV lytic infection, we assessed SMAD3 transcript levels after infection with WT and ∆miR-UL22A virus. As shown in Fig 2A, we observed a significant decrease in SMAD3 transcript levels upon infection with WT HCMV at 3 (p=0.05) and 6 (p<0.01) days post-infection (dpi), suggesting that the decrease in total and phosphorylated protein (Fig. 1C) is due, at least in part, to a decrease in transcription of SMAD3. Moreover, we observed only a partial restoration in SMAD3 transcript levels after ∆miR-UL22A infection, although significantly increased (p=0.02) compared to WT-infected cells at 6 dpi. We next treated mock, WT or ∆miR-UL22A virus-infected fibroblasts with TGFβ and assessed phospho-and total SMAD3 protein levels (Fig 2B). Cells infected with the miR-UL22A knockout virus showed enhanced SMAD3 phosphorylation and increased SMAD3 protein levels compared to WT-infected cells but did not reach the levels observed in mock-infected cells. We also assessed downstream transcriptional targets of TGFβ signaling after infection with WT or miR-UL22A knockout virus (Fig 2C). Infection with miR-UL22A knockout virus resulted in significantly increased JunB (p=0.03) and SERPINE (p<0.01) transcript levels compared to WT infected cells but again infection with the ∆miR-UL22A mutant virus did not restore transcript levels to that observed in mock-infected cells. Similar observations were made with endothelial cells (Fig 2D-E) infected with WT and miR-UL22A knockout viruses. These data indicate that miR-UL22A contributes to the blockade in TGFβ signaling observed during HCMV infection but, unlike in CD34^+^HPCs where mutation of miR-UL22A completely restores TGFβ signaling, other viral factors contribute to the reduction in SMAD3 transcript and protein levels during lytic infection.

**Fig 2.**
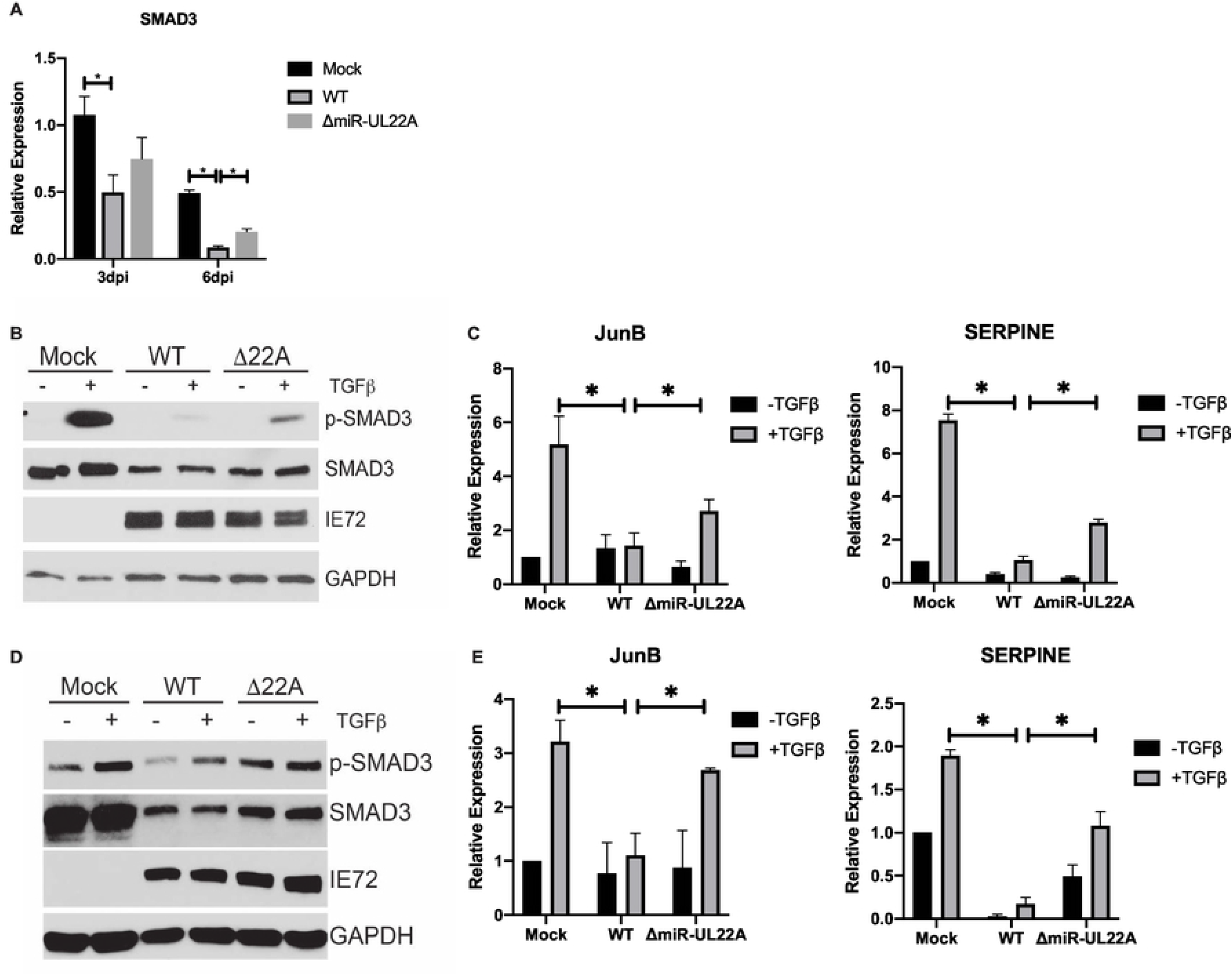
miR-UL22A targets SMAD3 during lytic infection. (A) NHDF were infected for 3 or 6 days with the indicated virus and then RNA was isolated and subjected to qRT-PCR for SMAD3. Experiments were performed in triplicate. NHDF (B) or hAEC (D) were infected with the indicated viruses for 48 hours followed by overnight serum starvation and stimulation with TGFβ (100pg/mL) for 4 hours. Protein lysates were subjected to immunoblotting using the indicated antibodies. NHDF (C) or hAEC (E) were infected as in (B) and RNA was isolated followed by qRT-PCR for JunB or SERPINE. Experiments were performed in triplicate. * p<0.05 by two tailed Student’s t test.

### miR-UL22A downregulation of SMAD3 expression is important for viral replication in the presence of exogenous TGFβ

The data presented here indicate that HCMV encodes multiple mechanisms to reduce SMAD3 expression and block canonical TGFβ signaling suggesting an anti-viral role for TGFβ during lytic infection. With this in mind, we assessed the functional consequences of TGFβ signaling on viral replication using the ∆miR-UL22A mutant virus which does not fully attenuate signaling through the canonical TGFβ signaling pathway (Fig 2). We also utilized a ∆miR-UL22A mutant that expresses a SMAD3 shRNA in place of the miR-UL22A hairpin (∆miR-UL22A/SMAD3shRNA). We have previously determined that expression of a SMAD3 shRNA in place of miR-UL22A results in SMAD3 protein levels similar to WT infection (9). Additionally, we observed that expression of an shRNA from the miR-UL22A locus restores TGFβ-responsive transcript levels to those observed during WT lytic infection (Fig 3A). We next performed multi-step growth analysis in fibroblasts infected with WT, ∆miR-UL22A and ∆miR-UL22A/SMAD3shRNA where exogenous TGFβ was added every 3 days. As shown in Fig 3B, replication of WT virus is unaffected by the addition of exogenous TGFβ. However, the miR-UL22A knockout virus, which shows an ∼1 log growth defect upon low multiplicity infection, was further inhibited for growth in the presence of exogenous TGFβ. Expression of a SMAD3 shRNA in place of miR-UL22A restored growth of the mutant virus to WT levels and showed no growth defect upon TGFβ treatment. This data suggests that reducing SMAD3 protein levels and blocking the canonical TGFβ signaling pathway is important for efficient lytic viral replication.

**Fig 3.**
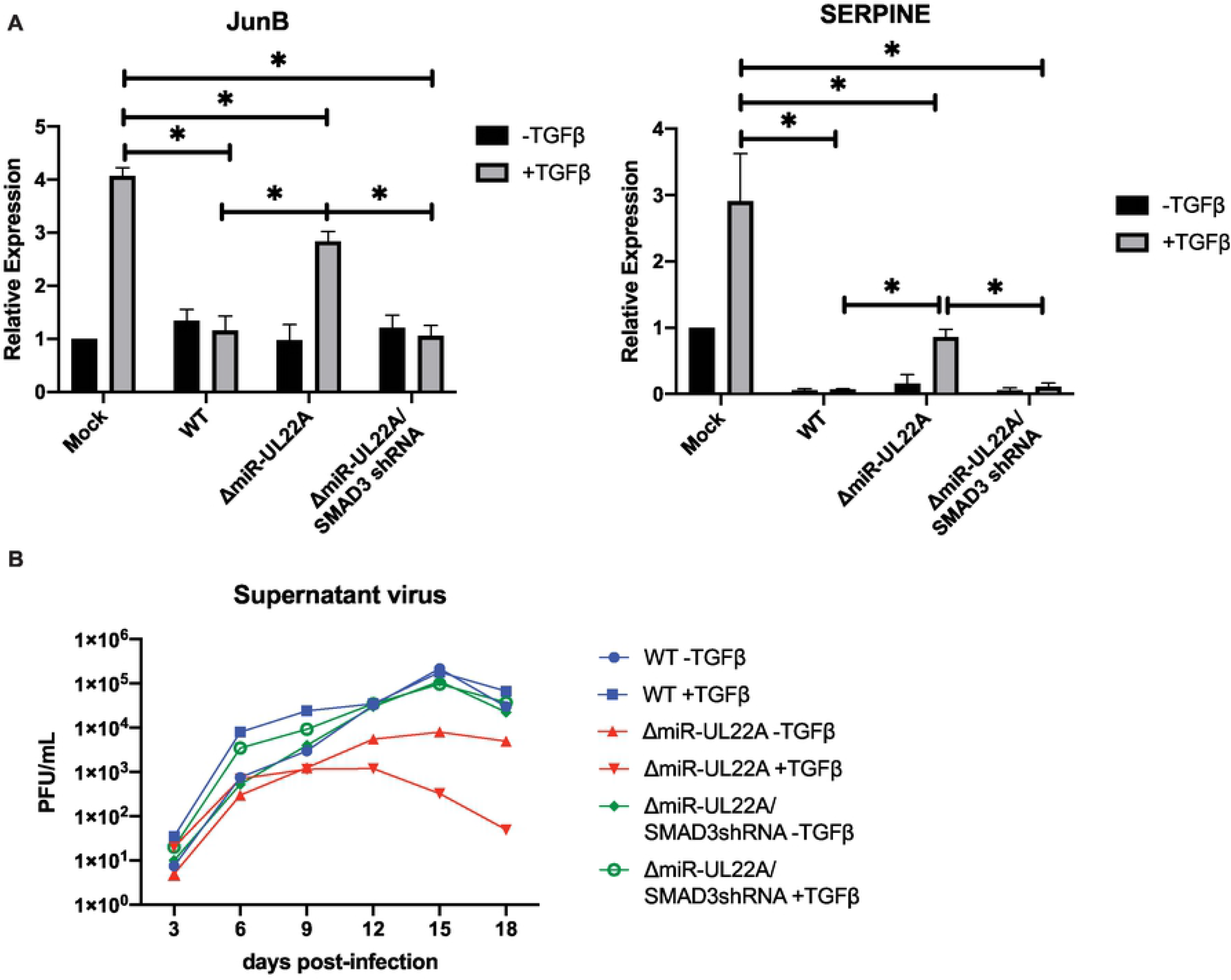
Targeting SMAD3 for downregulation is necessary for efficient lytic replication. (A) NHDF were infected with the indicated viruses for 48 hours followed by overnight serum starvation and stimulation with recombinant TGFβ (100pg/mL) for 4 hours. RNA was isolated and subjected to qRT-PCR for JunB and SERPINE. Experiments were performed in triplicate. * p<0.05 by two tailed Student’s t test. (B) NHDF were infected at 0.01 PFU/mL for 2 hours. 100pg/mL TGFβ was added immediately after infection and every 3 days throughout the experiment. Samples were harvested at the indicated timepoints and titered on NHDF. Experiment was performed in duplicate.

### miR-UL22A targeting of SMAD3 regulates expression of IFNβ and interferon stimulated genes

The negative effect of exogenous TGFβ on lytic replication of the ∆miR-UL22A virus and the restoration of mutant virus growth upon downregulation of SMAD3 implicates SMAD3 protein in anti-viral responses. SMAD proteins themselves have very weak affinity for SMAD binding elements in the promoters of targeted genes, and most often interact with additional DNA binding cofactors to mediate their effects (11). One cofactor known to interact with SMAD3 is interferon regulatory factor 7 (IRF7) which, along with SMAD3, has been shown to mediate activation of the IFNβ promoter (25). Thus, we asked whether miR-UL22A, through its ability to target SMAD3, could affect the induction of IFNβ and ISGs in response to a variety of stimuli. As shown in Fig 4, transfection of miR-UL22A, SMAD3 siRNA or IRF7 siRNA significantly blocked the induction of IFNβ or the ISG RSAD2 (Viperin) in response to the cytosolic DNA stimuli ISD90 or UV-HCMV (Fig 4A, C) and RNA stimuli, including the single-stranded RNA virus Sendai virus and polyI:C (Fig 4B, D). These data implicate both SMAD3 and IRF7 in IFNβ and ISG induction in fibroblasts and show that miR-UL22A, likely through targeting SMAD3, also interferes with IFNβ and ISG induction.

**Fig 4.**
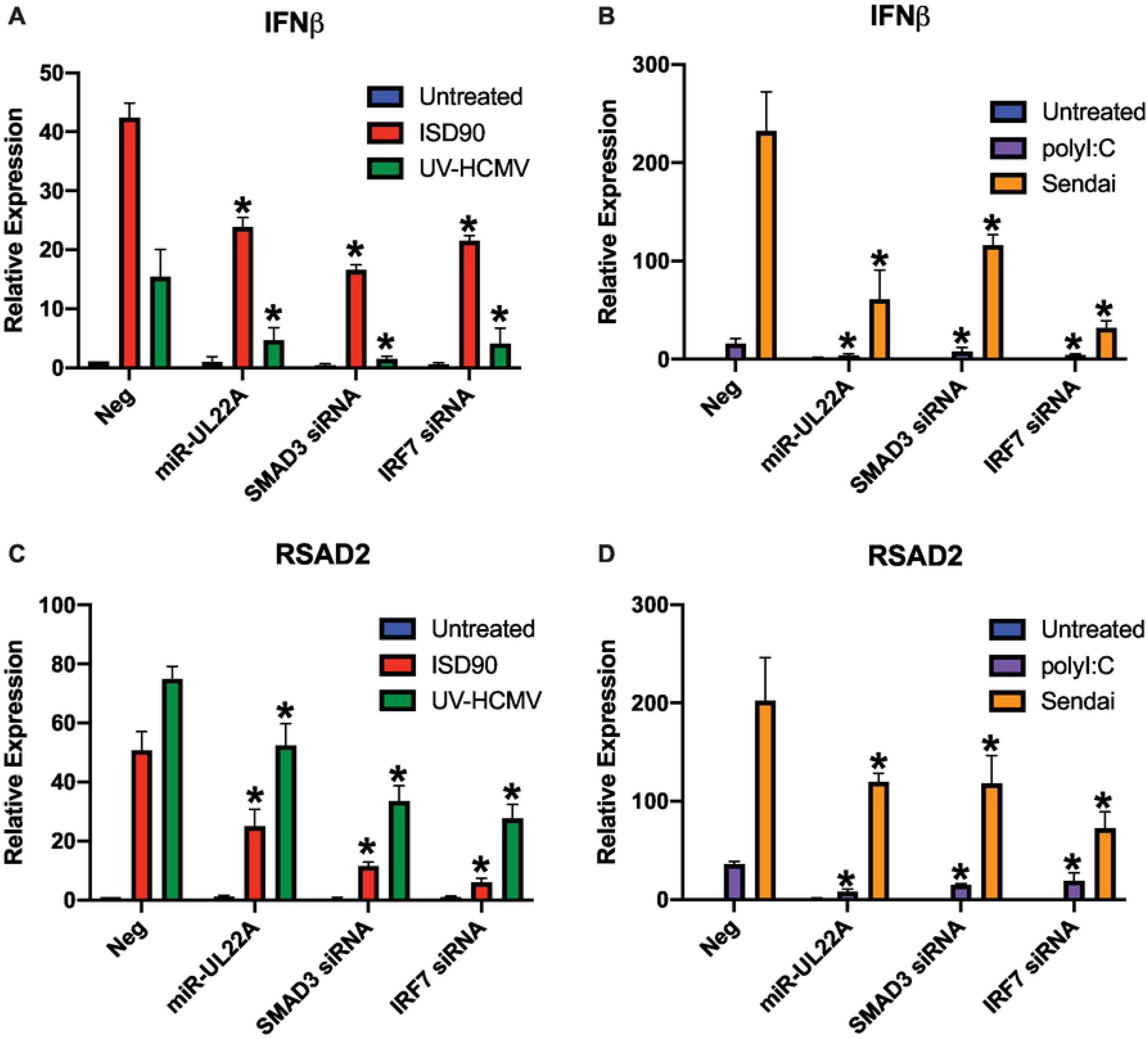
miR-UL22A and SMAD3 siRNA affect IFN and ISG induction in response to multiple stimuli. (A-D) NHDF were transfected with negative control, miR-UL22A mimic or SMAD3 or IRF7 siRNA. 24 hours post-transfection cells were either: (A, C) infected with UV-inactivated HCMV or transfected with ISD90 for a further 24 hours or (B, D) infected with Sendai virus or transfected with polyI:C for a further 24 hours. After this time, RNA was isolated and qRT-PCR was performed for IFNβ or RSAD2. All experiments were performed in triplicate. * p<0.05 by two tailed Student’s t test.

We next wanted to validate that the effects of miR-UL22A and SMAD3 knockdown on IFN and ISG induction occurred through the well characterized STING and JAK/STAT mediated signaling pathways. To do this we tested the effects of miR-UL22A and SMAD3 or IRF7 siRNA expression on IFN and ISG induction in previously constructed telomerized human fibroblast (tHF) cell lines deficient for STING or the type I IFN receptor IFNAR (∆STING and ∆IFNAR) (26-28). STING is a key regulator in innate immune signaling downstream of cGAS recognition of incoming viral DNA including the response to HCMV infection (29, 30). In WT tHF cells, expression of miR-UL22A, SMAD3 or IRF7 siRNAs reduced IFNα, IFNβ and RSAD2 induction in response to ISD90, in agreement with the data presented in Fig 4 in NHDF. In the absence of STING, IFNs and RSAD2 were not induced by ISD90 stimulation and miR-UL22A and SMAD3 or IRF7 siRNAs did not further affect expression (Fig 5A-C), indicating that SMAD3, as a target of miR-UL22A, functions downstream of STING-mediated innate immune signaling. In ∆IFNAR cells, the initial induction of IFNβ detected, but further amplification mediated by the IFNAR/JAK/STAT signaling pathway does not ensue (Fig 5D). We observed that miR-UL22A, SMAD3 and IRF7 siRNAs attenuated the (reduced) induction of IFNβ in ∆IFNAR cells (Fig 5D), indicating that they function in the initial induction of IFNβ that occurs prior to signal amplification through the JAK/STAT pathway. IFNα and RSAD2 expression was abrogated in ∆IFNAR cells and not further affected by expression of miR-U22A or SMAD3 or IRF7 siRNAs (Fig 5E-F) consistent with their functions as ISGs induced by JAK/STAT signaling, in the case of RSAD2, and directly dependent on IRF7 expression, in the case of IFNα (31-33). Thus, this data supports the hypothesis that miR-UL22A modulates SMAD3-mediated induction of IFNβ and downstream ISG induction.

**Fig 5.**
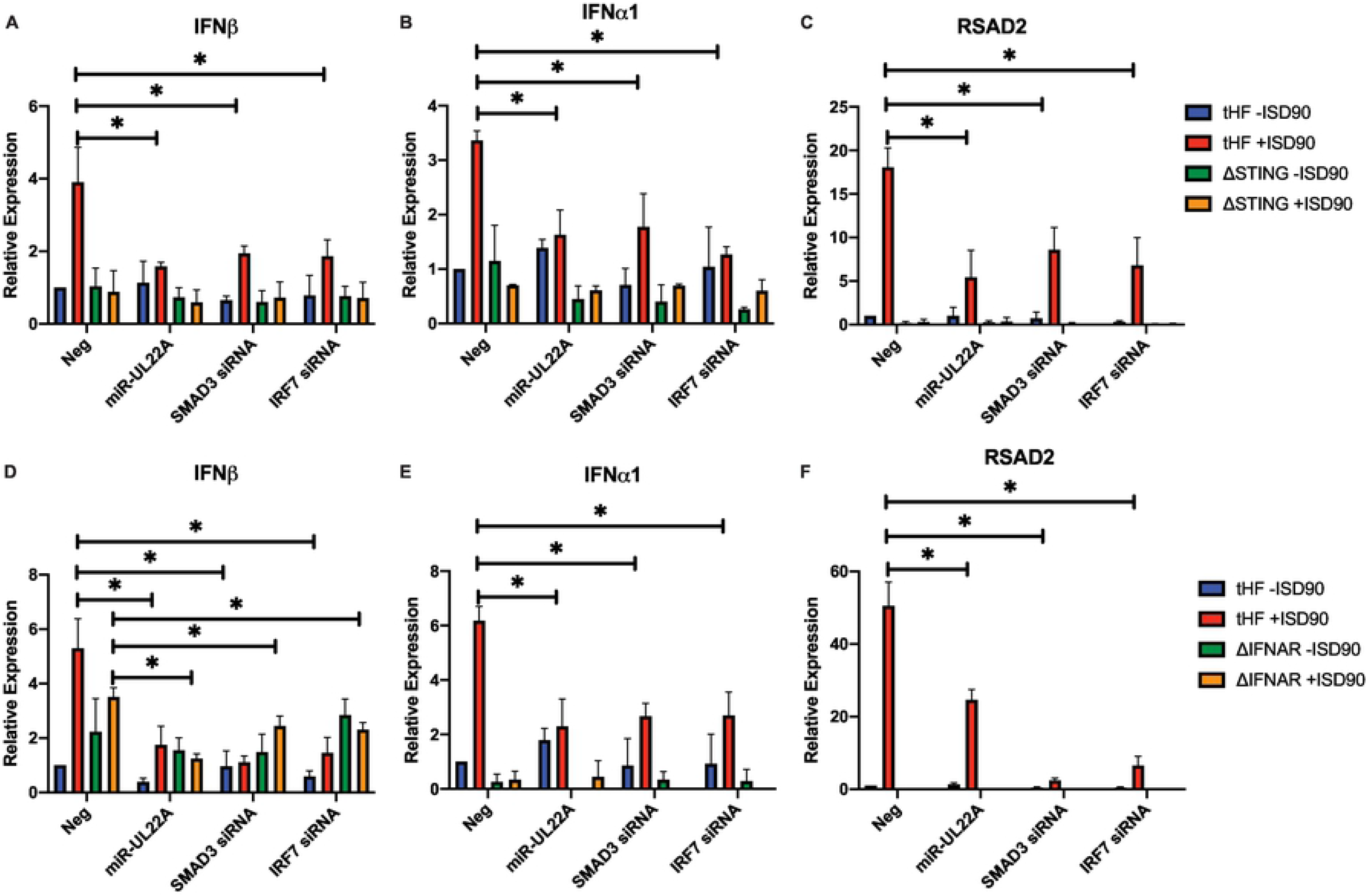
SMAD3 participates in STING-and IFN receptor-mediated signaling. tHF, ∆STING (A-C) or ∆IFNAR (D-F) tHFs were transfected with negative control, miR-UL22A mimic or SMAD3 or IRF7 siRNA for 48 hours after which cells were additionally mock transfected or transfected with ISD90 for 16 hours. RNA was isolated and subjected to qRT-PCR for IFNβ, IFNα1 or RSAD2. All experiments were performed in triplicate. * p<0.05 by two tailed Student’s t test.

In order to determine if the effect of miR-UL22A expression and knockdown of SMAD3 on IFN and ISG induction required IRF7, we derived an IRF7 knockout tHF cell line using CRISPR/Cas9 genome editing (27, 28, 34). Endogenous IRF7 protein levels are generally undetectable in many cell types, but its expression can be induced by IFN or other innate immune stimuli (33). As shown in Fig 6A, IRF7 expression is induced by transfection of the parental tHF cells with ISD90, but IRF7 protein was undetectable in the IRF7 knockout tHF cell line. In addition, IRF7 plays a key role in the induction of IFNα (33, 35), which we show is not induced in the IRF7 knockout cells following ISD90 treatment (Fig 6C), further validating the ∆IRF7 cell line. We next tested the effects of miR-UL22A and SMAD3 or IRF7 siRNA expression on IFN and ISG induction after treatment of ∆IRF7 tHF cells with ISD90. In ∆IRF7 cells, type I IFN and ISG induction is abrogated, and miR-UL22A or siRNAs targeting SMAD3 and IRF7 had no additional effect on gene expression consistent with the hypothesis that miR-UL22A, through targeting SMAD3, affects IRF7-mediated induction of IFNβ and downstream ISGs.

**Fig 6.**
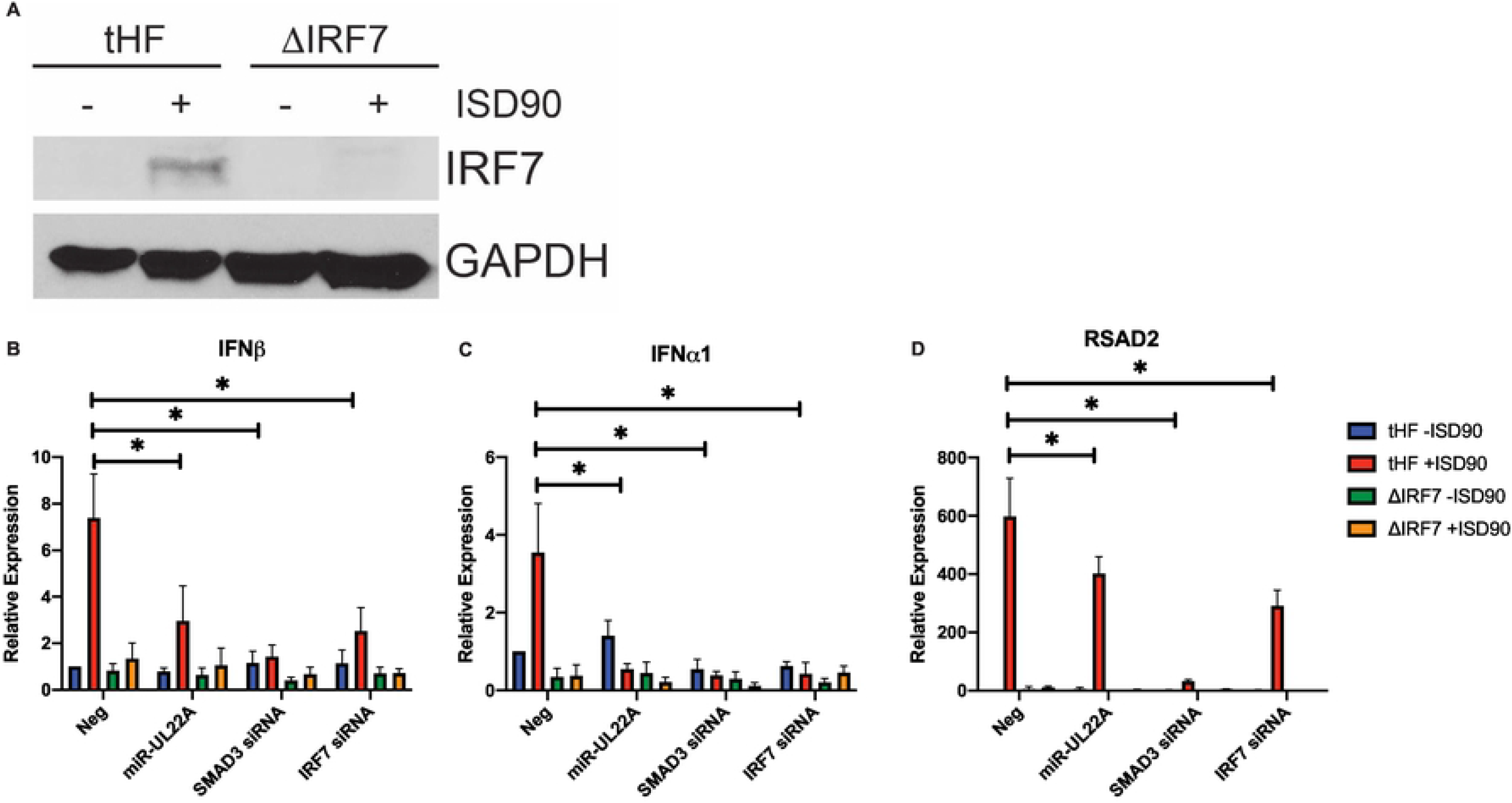
SMAD3 participates in IRF7-mediated signaling. (A) tHF or ∆IRF7 tHFs were transfected with ISD90 for 16 hours. Protein lysates were subjected to immunoblotting for IRF7 or GAPDH. (B-D) tHF or ∆IRF7 tHFs were transfected with negative control, miR-UL22A mimic or SMAD3 or IRF7 siRNA for 48 hours after which cells were additionally mock transfected or transfected with ISD90 for 16 hours. RNA was isolated and subjected to qRT-PCR for IFNβ, IFNα1 or RSAD2. All experiments were performed in triplicate. * p<0.05 by two tailed Student’s t test.

### miR-UL22A targeting of SMAD3 limits induction of type I interferons and ISGs during lytic infection

In order to assess the role of miR-UL22A regulation of SMAD3 and IRF7-mediated signaling in the context of HCMV infection, we next tested the induction of IFN transcripts following infection with WT and ∆miR-UL22A mutant viruses. As shown in Fig 7A and B, infection with the ∆miR-UL22A virus resulted in increased IFNβ and IFNα transcript accumulation both in the absence and presence of IFN treatement compared WT-and ∆miR-UL22A/SMAD3shRNA-infected cells. We also measured IFN secretion following infection with WT and miR-UL22A mutant virus (Fig 7C and D) and showed enhanced secretion of IFNβ and IFNα upon ∆miR-UL22A mutant virus infection that is reduced in cells infected with ∆miR-UL22A/SMAD3shRNA virus. We then assessed the induction of additional ISGs following infection with the ∆miR-UL22A mutant viruses and treatment with IFN. As shown in Fig 7E and F, induction of IRF7 and RSAD2 is enhanced in cells infected with ∆miR-UL22A mutant virus compared to WT-and ∆miR-UL22A/SMAD3shRNA-infected cells after IFN treatment but does not reach levels observed in IFN-treated, mock-infected cells. These data indicate that miR-UL22A, through targeting SMAD3, is important for reducing type I IFN and ISG production during lytic HCMV infection but also that other gene products are likely also involved in SMAD3 and IFN regulation.

**Fig 7.**
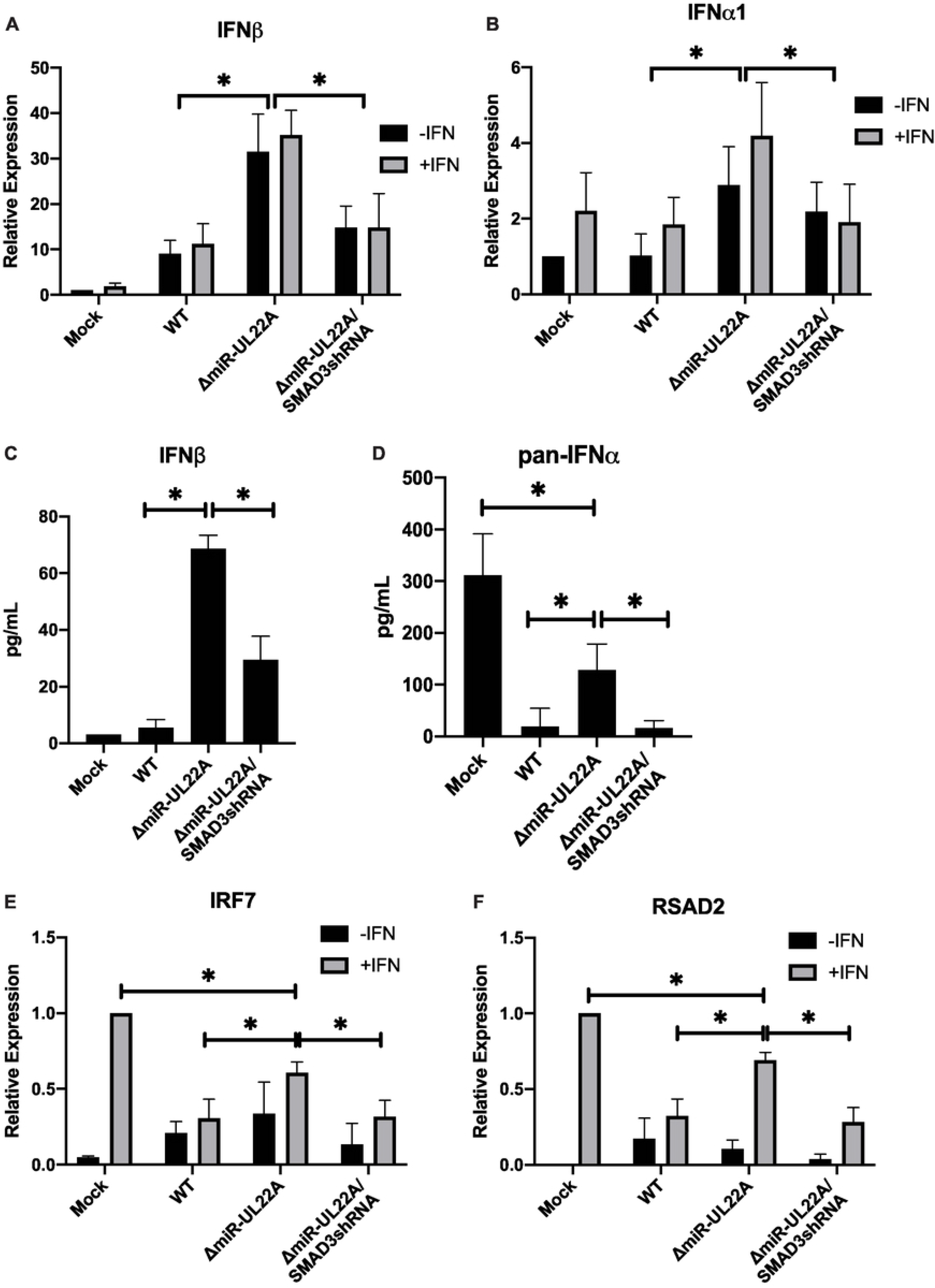
miR-UL22A targeting of SMAD3 regulates IFN and ISG induction. (A-B) NHDFs were infected with the indicated viruses for 48 hours followed by treatment with uIFN (1000U/mL) for 16 hours. RNA was isolated and subjected to qRT-PCR for IFNβ and IFNα1. (C-D) NHDFs were infected as in (A) and supernatants were harvested after 72 hours. ELISAs were performed for (C) IFNβ and (D) pan-IFNα. (E-F) NHDF were infected as in (A) and treated with IFN after 24 hours of infection. RNA was isolated 48 hours post-infection and subjected to qRT-PCR for IRF7 and RSAD2. All experiments were performed in triplicate. * p<0.05 by two tailed Student’s t test.

### TGFβ-mediated attenuation of lytic infection is mediated through SMAD3 and IRF7

Finally, in order to determine whether the TGFβ-mediated attenuation of replication of the ∆miR-UL22A virus is due to the cross-talk between TGFβ and innate immune signaling, we analyzed the growth of WT and miR-UL22A mutant viruses in WT and ∆STING or ∆IRF7 tHFs. Fig 8 demonstrates that while the ∆miR-UL22A mutant virus shows reduced virus released into the supernatant upon TGFβ treatement in parental tHFs compared to WT and ∆miR-UL22A/SMAD3shRNA viruses (Fig 8A), this defect was abrogated in cells lacking STING (Fig 8B), further supporting the role of innate immune signaling in attenuating replication of a ∆miR-UL22A mutant virus. Moreover, the defect in ∆miR-UL22A mutant virus growth is also abrogated in ∆IRF7 cells, (Fig 8C) indicating that IRF7 is directly involved in impeding ∆miR-UL22A growth in response to TGFβ. Together, these data support the hypothesis that SMAD3 and IRF7 cooperate to induce IFN production during HCMV infection, which has negative effects on viral replication.

**Fig 8.**
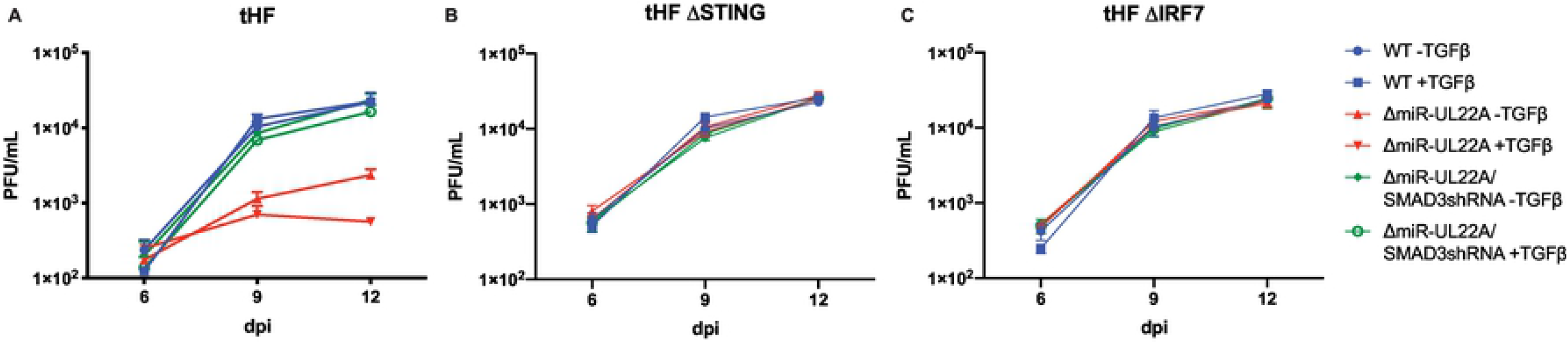
TGFβ impairs lytic replication in coordination with IRF7. tHF (A), ∆STING (B) or ∆IRF7 (C) tHFs were infected at 0.01 PFU/mL. TGFβ was added (100pg/mL) after initial infection and again at days 6 and 9. Supernatants were harvested and titered in NHDF. Experiments were performed in duplicate.

## Discussion

In this study we show that SMAD3, a TGFβ receptor-associated SMAD, cooperates with IRF7 to induce type I IFN during HCMV infection. miR-UL22A, through downregulating SMAD3 expression, plays an important role in regulating IRF7-mediated IFN production and viral replication during lytic infection. Infection with virus lacking miR-UL22A results in enhanced IFN production and release, enhanced downstream ISG induction and inhibition of growth in the presence of exogenous TGFβ. However, if the ∆miR-UL22A mutant expresses an shRNA targeting SMAD3, type I IFN and ISG induction as well as viral replication returns to WT levels, directly implicating the regulation of SMAD3 expression by miR-UL22A in modulation of IFN production and efficient viral replication.

Herpesviruses manipulate the intrinsic and innate IFN signaling pathways to aid in their replication cycles, which is especially important during lytic infection, where efficient viral gene expression and new virion production is paramount. In order to induce the production of IFNs after recognition of viral infection by pattern recognition receptors, the transcription factors IRF3 and IRF7 are phosphorylated by the kinase TBK1, translocate to the nucleus and, along with c-Jun, ATF2, NFκB and CBP/p300, bind to the *Ifnb* promoter to induce its expression (36). Autocrine IFN signaling through the IFN receptor then stimulates the production of more IRF7 and thus more type I IFN via positive feedback (35, 37), establishing IRF7 as a ‘master regulator’ of IFN production (35). In this study we show that SMAD3, a TGFβ receptor-associated SMAD, cooperates with IRF7 to induce type I IFNs during HCMV infection, highlighting a unique interconnection between TGFβ and IFN signaling.

An interaction between IRF7 and SMAD3 was first postulated due to the similarity of each transactivation domain and the fact that, upon phosphorylation, both proteins undergo structual rearrangements that promote complex formation (25). SMAD3 was also shown to interact with IRF7 (but not IRF3) at the *Ifnb* promoter in mouse embryonic fibroblasts and this interaction was critical for *Ifnb* transcription (25). Here we show that both IRF7 and SMAD3 are required for induction of IFN and ISG transcripts in response to a variety of PAMPs in human fibroblasts (Fig 4). The effect of miR-UL22A and SMAD3 siRNA on type I IFN and ISG induction was abrogated in the absence of the signaling adaptor STING (Fig 5A-C), indicating that the SMAD3, like IRF7, functions as a component of the IFN-terminal innate immune signaling pathway. The hypothesis that SMAD3 and IRF7 work together in the initial induction of IFNβ is supported by the observations using ∆IFNAR cells, which are capable of the initial IFNβ production following infection but cannot amplify the response. Expression of miR-UL22A and SMAD3 or IRF7 siRNA blocks the initial induction of IFNβ in ∆IFNAR cells (Fig 5D). Furthermore, miR-UL22A and SMAD3 siRNA have no effect on the low levels of IFNβ transcription in IRF7 knockout cells (Fig 6B-D), suggesting a cooperative function of SMAD3 and IRF7 in the induction of type I IFNs and downstream ISG induction in human fibroblasts.

The functional significance of SMAD3 targeting by miR-UL22A during lytic infection is underscored by the increased IFNα and IFNβ expression and secretion upon infection with a ∆miR-UL22A mutant virus (Fig 7). This enhanced type I IFN response results in increased ISG induction in the absence of miR-UL22A expression, although not to levels observed in mock-infected cells indicating that likely other SMAD3-targeting gene products are involved in this process (Fig 7 E&F). Critically, replacing the miR-UL22A hairpin locus with an shRNA targeting SMAD3 reduces type I IFN expression and secretion, along with ISG induction, to levels seen during WT infection. This indicates both the necessity and sufficiency of targeting SMAD3 for HCMV-mediated impairment of IFN. The negative effect of TGFβ on replication of the ∆miR-UL22A mutant virus is abrogated in cell lines lacking STING or IRF7 (Fig 8), further supporting the hypothesis that SMAD3 and IRF7 function together to limit replication of HCMV through the induction of a type I IFN response.

Regulation of IRF7 expression and function is utilized by α− and γ-herpesviruses as a means to dampen the IFN response during infection. Kaposi’s Sarcoma-associated Herpesvirus (KSHV) and Herpes Simplex virus (HSV) encode proteins that sequester or degrade IRF7 (38-45). In contrast, during Epstein Barr virus (EBV) latent infection, IRF7 expression is stimulated by latent membrane protein-1 (LMP-1) (46-49) which in turn regulates expression of the EBNA1 Q promoter (50) and LMP-1 itself. However, during reactivation, EBV IE proteins BZLF1 (51) and LF2 (52) bind and repress IRF7 activity and BRLF1 downregulates IRF7 expression and IFNβ production (53). Thus, along with the data presented here for HCMV, it is clear that targeting IRF7 expression and/or function during lytic infection is a common theme amongst the herpesvirus family.

The role of TGFβ signaling in herpesvirus infection is highly relevant yet complex, with differing effects on lytic and latency stages of the lifecycle. TGFβ treatment of EBV latently infected cells can induce reactivation via SMAD binding to the *BZLF1* promoter (54). Thus, EBV encodes factors that directly or indirectly interfere with components of the TGFβ signaling pathway (55-57). Additionally, EBV proteins upregulate the cellular miRNA miR-146a which directly targets SMAD4 (58, 59). Likewise, KSHV proteins induce the expression of the cellular miR-17-92 family, which targets SMAD2 (60). KSHV also encodes proteins (61-63) and miRNAs (64, 65) that target components of the TGFβ signaling pathway. In contrast, infection of human mononuclear cells with HSV-1 induces TGFβ production (66) and use of conditional TGFβ knockout mouse models suggests that TGFβ signaling, while dampening the innate immune response, enhances HSV-1 latency (67).

Similar to HSV-1, the HCMV major immediate early proteins can activate the TGFβ promoter, which occurs indirectly through IE2 transactivation of the cellular immediate early transcription factor EGR-1, but was also shown to require additional viral factors during infection (6, 7). Recent studies implicate miR-US5-2 targeting of the transcription repressor NAB1 as one of the possible additional mechanisms (9). Furthermore, HCMV IE1 and IE2 induce the expression of MMP-2 in renal tubular epithelial cells, which may result in enhanced activation of latent TGFβ in the extracellular matrix of HCMV-infected cells and contribute to the fibrosis observed during transplantation (68). Thus, CMV infection induces the production of TGFβ by multiple mechanisms but encodes multiple factors to block the canonical TGFβ signaling pathway (Figs 1-3).

While removal of the pre-miR-UL22A sequence from the viral genome results in enhanced SMAD3 protein levels and expression of classical downstream TGFβ transcriptional targets during lytic infection, responses do not return to levels observed in mock-infected cells (Fig 2B-E, 7E&F), suggesting that the virus uses additional mechanisms (utilizing viral proteins and/or additional miRNAs or long non-coding RNAs) to alter SMAD3 mRNA and protein levels and inhibit the TGFβ signaling pathway. Possible additional mechanisms utilized by HCMV to to manipulate the TGFβ signaling pathway remain to be identified, but could include inhibiting SMAD3 transcription initiation, affecting the expression or stability of additional members of the signaling pathway or induction of negative pathway regulators, such as the I-SMADs. Interestingly, HCMV blocks the signaling pathway of the related TGFβ family member activin by directly targeting the activin receptor (ACVR1B) using miR-UL148D to prevent the production and release of IL6 in monocytes (69).

The interplay between type I IFN and TGFβ signaling during HCMV infection highlights the complex interconnection of signaling pathways that are regulated by viral proteins and non-coding RNAs. An inability to downregulate SMAD3 during latent infection results in a loss of viral genomes from the infected cells (9) and whether this phenotype is due to enhanced IFN signaling remains to be determined. Future studies will explore how this novel interplay between the TGFβ and IFN signaling pathways is important for HCMV latency in hematopoietic progenitor cells.

## Materials and Methods

### Cell lines

Normal human dermal fibroblasts (NHDF), human foreskin fibroblasts (stably transduced with constitutively expressed human telomerase reverse transcriptase and the IRF/IFN-responsive pGreenFire-ISRE lentivector; tHF) (27) and the tHF ∆STING (26), ∆IFNAR (28) and ∆IRF7 cell lines were cultured in Dulbecco’s modified Eagle’s medium (DMEM) supplemented with 5% heat-inactivated fetal bovine serum (FBS; HyClone), 100 units/ml penicillin, and 100ug/ml streptomycin (ThermoFisher). Human aortic endothelial cells (AEC) (CC-2535; Lonza) were cultured in EBM-2 basal medium with EGM-2 SingleQuots™ supplement excluding Heparin (Lonza), as well as 10% FBS, penicillin, and streptomycin. All cells were maintained at 37**°**C and 5% CO2. Recombinant human TGFβ and Universal Type I IFN was obtained from R&D Systems.

### HCMV Constructs and infections

HCMV used in this study include BAC-generated WT TB40/E expressing GFP from the SV40 promoter (70), a TB40/E mutant virus lacking the pre-miR-UL22A sequence or a TB40/E mutant virus with the pre-miR-UL22A sequence replaced by a SMAD3 shRNA generated by galK-mediated recombination (9). All virus stocks were propagated and titered on NHDFs. Fibroblasts were infected with HCMV at three plaque-forming units (PFU)/cell and hAEC were infected with HCMV at five PFU/cell for 2 hours at 37°C. After this time, the inoculum was removed and replaced with fresh medium and samples were harvested as appropriate for each experiment. For experiments involving TGFβ stimulation, cells were infected as above for 48 hours followed by serum starvation overnight. The next day cells were treated with 100pg/mL TGFβ for 4 hours. Multi-step growth curves were performed in duplicate using NHDF, tHF or derivitives infected with 0.01 PFU/cell in DMEM containing 1% FBS and recombinant TGFβ (100pg/mL) was added immediately after infection and every 3 days thereafter. UV inactivation of HCMV was performed using the Spectrolinker XL-1000 (Spectronics Corporation) by exposing virus resuspended in 200μl for 30s at 600 μJ three times sequentially (71). Sendai virus (SeV) was obtained from Charles River Laboratories and used at 160 hemagglutination units (HAU)/mL.

### Transfections

NHDF or tHF cells seeded in 12 well plates were transfected with 40uM siRNAs (SMAD3 and IRF7; ThermoFisherScientific) or miRNA mimics (custom designed; IDT) per well using Lipofectamine RNAiMax (ThermoFisherScientific) according to the manufacturer’s instructions. 33ug/mL ISD90 (IDT) and 16.5ug/mL polyI:C (Invivogen) was transfected into cells using Lipofectamine 3000 according to the manufacturer’s instructions.

### qRT-PCR

Total RNA was isolated from transfected or infected cells using the Trizol RNA isolation method. cDNA was prepared using 1000ng of total RNA and random hexamer primers. Samples were incubated at 16°C for 30 minutes, 42°C for 30 minutes and 85°C for 5 minutes. Real-time PCR (Taqman) was used to analyze cDNA levels in transfected or infected samples. An ABI StepOnePlus Real Time PCR machine was used with the following program for 40 cycles: 95°C for 15 sec and 60°C for one minute. Relative expression was determined using the ∆∆Ct method using 18S as the standard control. JunB, SERPINE, SMAD3, TGFB1, IFNA1, IFNB1, RSAD2, IRF7 and 18S primer/probe sets were obtained from ThermoFisher Scientific.

### Immunoblotting

Protein extracts were run on an 8% SDS-PAGE, transferred to Immobilon-P Transfer Membranes (Milipore Corp., Bedford, MA), and visualized with specific antibodies: phosho-SMAD3 (Abcam), total SMAD3 (Abcam), IE86 (mAb 810; Millipore), IRF7 (Santa Cruz) and GAPDH (Abcam). Relative intensity of bands detected by western blotting was quantitated using ImageJ software.

### ELISAs

Supernatants harvested from infected cells were centrifuged at maximum speed for 30sec to remove cell debris and stored at −80°C prior to cytokine measurements. IFNα was quantified using the pan-IFNα ELISA kit (Stem Cell Technologies). IFNβ was quantified using human IFN-beta Quantikine ELISA kit (R&D Systems). All measurements were made following the manufacturers’ protocols.

### Construction of IRF7 KO fibroblasts

Genome editing using lentivirus-mediated delivery of CRISPR-Cas9 components was performed generally as described previously (27, 28, 34). Breifly, a 20 nucleotide guide RNA (gRNA) sequence targeting the IRF7 protein-coding region was inserted into the lentiCRISPRv2 vector (Addgene; catalog #52961). The IRF7 guide sequences used was: TACACCTTGTGCGGGTCGGC. tHF cells stably transduced with the IFN-responsive pGreenFire-ISRE lentivector (System Biosciences) were further stably transduced with the lentiCRISPRv2 vector, selected using puromycin at 3ug/mL, and IRF7 knockdown was confirmed by western blotting.

### Statistical Analysis

The Student’s two tailed t test (Microsoft Excel software) was used to determine p values. Results were considered significant at a probability (p) < 0.05.

## Acknowledgments

The authors would like to acknowledge members of the Nelson and Defilippis labs for helpful discussions.

